# Human untargeted metabolomics in the gut microbiome era: ethanol vs methanol

**DOI:** 10.1101/2024.09.23.614605

**Authors:** Simone Zuffa, Vincent Charron-Lamoureux, Caitriona Brennan, Madison Ambre, Rob Knight, Pieter C. Dorrestein

## Abstract

Untargeted metabolomics is frequently performed on human fecal samples in conjunction with sequencing to unravel gut microbiome functionality. As sample collection efforts rapidly expand, with individuals often collecting specimens at home, metabolomics experiments should adapt to accommodate the safety and needs of bulk off-site collections. Here, we show that a 95% ethanol, safe to be shipped and handled, extraction pipeline recovers comparable amounts of metabolites as a validated 50% methanol extraction, preserving metabolic profiles differences between subjects. Additionally, the fecal metabolome remains relatively stable when stored in 95% ethanol for up to a week at room temperature. Finally, we suggest a metabolomics data analysis workflow using robust centered log ratio transformation, which removes variance introduced by different sample weights, allowing for reliable and integration-ready untargeted metabolomics experiments in gut microbiome studies.

## Introduction

Humans are colonized at birth by microorganisms, forming complex communities known as microbiota, which evolve through the course of the host life, shaping and influencing its physiology and metabolism^1^. Among others, the gut microbiota has been shown to actively influence neurodevelopment before^2,3^ and after *partum*^*4–6*^, and to induce distal tumors via carcinogenic metabolism^7^. For these reasons, elucidating the functionality of these microbial communities has become paramount and mass spectrometry-based untargeted metabolomics has established itself as the to-go tool for these types of investigations^8^.

Multiomics large-scale longitudinal or cross-sectional microbiome studies represents a challenge both for sequencing and metabolomics, as samples are usually collected in different settings and cannot always be immediately snap-frozen in liquid nitrogen and stored at -80 °C, which is considered the gold standard. Additionally, when designing these large-scale studies safety of subjects and shipping costs, which dramatically increase if refrigeration is involved, should be taken into consideration. The microbiome field tackled these problems by showcasing that fecal microbiome collection and storage in 95% ethanol (EtOH) at room temperature (RT) stabilizes the microbial communities up to 8 weeks^9^. Most importantly, EtOH is also safer to handle when compared to other alcohols like methanol (MeOH), which is extremely toxic and requires special equipment to be properly handled. Building on this, we recently introduced the Matrix Method^10^, which employs a high-throughput pipeline that leverages sample collection in single barcoded Matrix™ tubes containing 95% EtOH and automatized robots. This method not only reduces costs, time, and well-to-well DNA contamination but enables the extraction of metabolites from the same sample, offering an all-in-one solution for unlocking a streamlined multi-omics analysis. EtOH extraction from fecal metabolomics studies has been tested before^11–14^ but an in-depth downstream data analysis comparison with another relevant extraction method (50% MeOH)^15–18^, that was used for the discovery of novel bile acids conjugates^19^ and the recently introduced reverse metabolomics approach^20^, is missing.

Here we showcase that the 95% EtOH extraction, part of the Matrix Method bulk microbiome analysis, yields equivalent results of a standard, but more laborious and time consuming, 50% MeOH extraction for human fecal samples. We also show that EtOH can stabilize the fecal metabolome at RT for up to a week, allowing for safe off-site collection and reduced shipping costs. Finally, we highlight how a robust center log ratio transformation data analysis workflow for untargeted metabolomics data overcomes problems of uneven sampling collection and allows for better integration with microbiome data.

### Experimental Section

#### Sample Collection and Extraction

Human fecal samples were collected and immediately stored at -80 °C from volunteers under approved protocols from the University of California San Diego (IRB#141853) and informed consent. Three fecal samples from three different subjects were randomly selected and aliquoted for untargeted metabolomics analysis. Multiple aliquots of different weights (10 mg, 20 mg, and 30 mg) were generated in triplicates from the three different fecal samples. Aliquots were either transferred in 95% (v/v) ethanol (EtOH) or 50% (v/v) methanol (MeOH). Samples were immediately extracted, except for a subset of samples that were left in 95% EtOH for either 24 h or 1 week at RT to check fecal metabolome stability. Samples to which 400 µL of 95% EtOH was added were extracted via the Matrix Method pipeline^10^, simply consisting in shaking samples at 1,200 rpm for 2 min in a SpexMiniG plate shaker (SPEX SamplePrep part #1600, NJ, USA), followed by a 5 min centrifugation step at 2,700 g. Supernatant (400 µL) was then collected and stored at -80 °C for downstream analysis. Samples to which 800 µL of MeOH was added underwent a validated extraction protocol^20^, involving homogenization with a 5 mm stainless steel bead in a TissueLyser II (QIAGEN) for 5 min at 25 Hz, incubation at 4 °C for 30 min, and centrifugation at 21,130 g for 10 min. Supernatant (400 µL) was then collected and dried overnight using a CentriVap. Samples were then stored at -80 °C until resuspension. All supernatants, from EtOH and MeOH extractions, were dried overnight using a CentriVap and then resuspended in 200 µL of 50% MeOH containing 1 µM of sulfamethazine as internal standard. A pooled sample (QCpool) was then generated by collecting and mixing 10 µL from each biological sample and aliquoting 200 µL. Blank samples, consisting only of extraction solution, were also prepared. Finally, samples were incubated for 1 h at -20 °C, centrifuged at 21,130 g for 10 min, and transferred in 2 mL glass vial (Thermo Scientific) for ultra high performance liquid-chromatography tandem mass spectrometry (UHPLC-MS/MS) analysis.

#### UHPLC-MS/MS Experiment

Samples were randomized and analyzed using an untargeted metabolomics analysis platform comprising a Vanquish UHPLC (ultra-high performance liquid chromatography) system coupled to a Q-Exactive Orbitrap mass spectrometer (Thermo Fisher Scientific). The chromatography system consisted in a Phenomenex C18 column (1.7 µm particle size, 2.1 mm x 50 mm) and a mobile phase of solvent A (water + 0.1% formic acid) and solvent B (acetonitrile + 0.1% formic acid). Injections of 5 μL of samples, with a flow rate of 0.5 mL/min, followed this gradient: 0-1 min 5% B, 1-7 min 5-99% B, 7-8 min 99% B, 8-8.5 min 99-5% B, 8.5-10 min 5% B. MS/MS data was acquired in data-dependent acquisition (DDA) mode using positive electrospray ionization (ESI+). Briefly, ESI parameters were set as following: 53 L/min sheath gas flow, 14 L/min aux gas flow rate, 3 L/min sweep gas flow, 3.5 kV spray voltage, 269°C intel capillary, and aux gas heater set to 438°C. MS scan range was set to 100 – 1500 *m/z* with a resolution at *m/z* 200 set to 35,000 with 1 microscans. Automatic gain control (AGC) was set to 5E4 with a maximum injection time of 50 ms. Up to 5 MS/MS (TopN = 5) spectra per MS1 were collected with a resolution at *m/z* 200 set to 17,500 with 1 microscans. Injection time was 50 ms with an AGC target of 5E4. The isolation window was set to 2.0 *m/z*. Normalized collision energy was set to a stepwise increase of 20, 30, and 40 eV with an apex trigger set to 2-15 s and a dynamic exclusion of 10 s.

#### UHPLC-MS/MS Data Processing

Obtained .raw files were converted into .mzML open-access format using ProteoWizard MSConvert^21^ and deposited on GNPS/MassIVE under the accession number MSV000095260. Feature detection and extraction was performed via MZmine 3.9 via batch processing^22^. The .xml file used for batch processing can be found in the associated GitHub page. Briefly, data was imported using MS1and MS2 detector via factor of lowest signal with noise factors set to 3 and 1.1 respectively. Sequentially, mass detection was performed and only ions acquired between 0.5 and 8 min, with MS1 and MS2 noise levels set to 5E4 and 1E3 respectively. Chromatogram builder parameters were set at 5 minimum consecutive scans, 1E5 minimum absolute height, and 10 ppm for *m/z* tolerance. Smoothing was applied before local minimum resolver, which had the following parameters: chromatographic threshold 85%, minimum search range retention time 0.2 min, minimum ratio of peak top/edge 1.7. Then, 13C isotope filter and isotope finder were applied. Features were aligned using join aligner with weight for *m/z* set to 3 and retention time tolerance set to 0.2 min. Features not detected in at least 3 samples were removed before performing peak finder. Ion identity networking and metaCorrelate were performed before exporting the final feature table. The GNPS and SIRIUS export functions were used to generate the feature table containing the peak areas and the .mgf files necessary for downstream analyses. Feature based molecular networking was performed in GNPS2 (https://gnps2.org/status?task=40d2affb3df544d4a2bbf6841a62d45a#)^23^. Molecular classes of ions with *m/z* < 800 were predicted via CANOPUS using SIRIUS 5.8^24^.

#### Data Analysis

Feature table was imported in R 4.2.2 for downstream data analysis. Total extracted peak area per sample was calculated and correlated to sample run order to identify possible acquisition problems during the run. The internal standard (IS) peak present in each sample (Sulfamethazine [M+H], *m/z* 279.0908 and retention time 2.26 minutes) was also extracted and correlated to the sample run order. The coefficients of variance (CV) of 6 different standards (Amitriptyline, Sulfadimethoxine, Sulfamethazine, Sulfamethizole, Sulfachloropyridazine, and Coumarin 314) present in the QCmix sample, which was run every 10 biological samples, were inspected to evaluate run quality. CVs of each acquired feature were also calculated using the QCpool^25^, which was also run every 10 biological samples. CV was calculated by dividing the mean of the extracted peak areas by the standard deviation. Feature table was cleaned via blank subtraction. Features only detected in the blank and QCmix samples or which mean peak areas were not at least 10 times the ones observed in the QCpool were discarded. The package ‘homologueDiscoverer v 0.0.0.9’ was used to remove detected PEGs (polyethylene glycol)^26^. Features with near zero variance were removed using ‘caret v 6.0’^27^. The package ‘mixOmics v 6.22’ was used for multivariate analysis^28^. Principal component analysis (PCA) and partial least square discriminant analysis (PLS-DA) were performed after robust center log ratio transformation (RCLR) via ‘vegan v 2.6’^29^. In the PCA models, PERMANOVA was used to evaluate group centroid separation while PERMDIPS2 was used to evaluate homogeneity of variance. PLS-DA models performances were evaluated using leave-one-out (loo) cross-validation. Variable importance (VIP) scores were calculated per feature and features with VIPs > 1 were considered significant. The package ‘UpSetR v 1.4’ was used to generate the upset plots^30^. High density regions (HDRs) plots were generated using ‘ggdensity v 1.0’. Log2 fold changes (Log2FC) were calculated by taking the log2 of divided means of the relative abundance of the peak areas of the group of interest. When the mean was 0 a pseudocount of 1E-9 was added. Linear mixed effect models were created using ‘lmerTest v 3.1’, using subject id as random effect. The packages ‘tidyverse v 2.0’ and ‘ggpubr v 0.6’ were used for data manipulation and visualization. Code for the analysis and to generate the figures is available on GitHub (https://github.com/simonezuffa/Manuscript_Matrix_Metabolomics).

## Results and Discussion

Fecal samples from three different individuals were aliquoted to generate 81 replicates (***Figure 1***). Triplicates of different weights, 10 mg, 20 mg, and 30 mg, were extracted via two different pipelines. The first one consisted of a 50% MeOH extraction protocol, previously described^20^, while the second one involved a 95% EtOH extraction protocol part of the recently introduced Matrix Method pipeline^10^. Two batches of triplicates collected in 95% EtOH were left at RT for one day and one week respectively to replicate possible collection scenarios of storing, shipping, and assessing fecal metabolome stability in EtOH.

**Figure 1.**
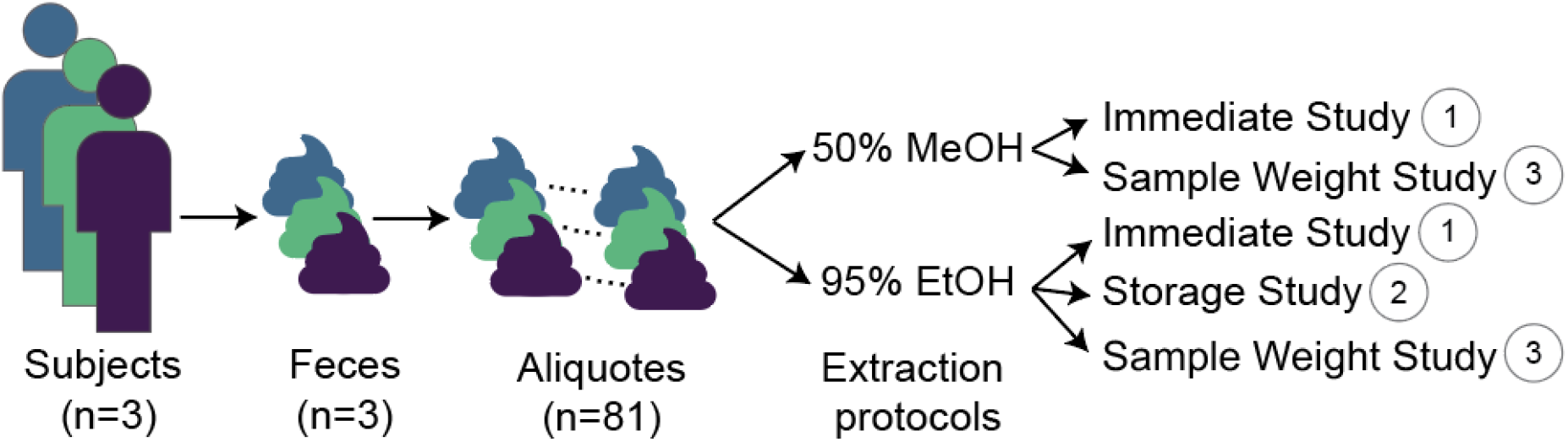
Study design. Fecal samples were collected from three different human subjects and aliquoted to investigate two different extraction methods: 95% ethanol (EtOH) and 50% methanol (MeOH). Three different studies were conducted. 1) Aliquoted 20 mg samples were immediately processed after thawing to investigate metabolites recovery differences between 50% MeOH and 95% EtOH. 2) Aliquoted 20 mg samples were immediately processed or left at room temperature for one day and one week in 95% EtOH to investigate fecal metabolome stability in EtOH. 3) Aliquoted 10 mg, 20 mg, and 30 mg were extracted via both 95% EtOH and 50% MeOH pipelines to investigate a weight-bias free data analysis workflow.

### 95% EtOH extraction recapitulates 50% MeOH extraction

Unsupervised dimensionality reduction via principal component analysis (PCA) of 20 mg triplicates showed clear clustering of subject fecal metabolic profiles using both 95% EtOH and 50% MeOH extractions (***Figure 2A***). PERMANOVA identified subject id as the highest source of variance in the data (R^2^ = 0.47, F = 10.98, p < 0.001), followed by extraction method (R^2^ = 0.14, F = 6.68, p < 0.001). Out of a total of 4,616 unique metabolic features obtained in this study, 75% (3,463) were recovered by both extraction methods (***Figure 2B***). These included 94% (235) of the total annotated features via the GNPS libraries. Interestingly, the EtOH extraction method appeared to capture a higher number of additional features (849) compared to the MeOH extraction (304). These extraction-specific features were classified as small peptides and fatty acid conjugates for MeOH and oligopeptides, glycerolipids, and fatty amides for EtOH, according to CANOPUS NPC superclass predictions (***Supplementary Table 1***). Subject pairwise PCA and PLS-DA models were generated for each extraction method to identify metabolic features responsible for subject discrimination. These features were then selected to compare both extraction methods. All PCA and PLS-DA score plots displayed clear clustering by subject (***Supplementary Figure 1***). All PLS-DA models obtained a classification error rate of 0, indicating a perfect discriminating performance. Extracted VIP scores from the PLS-DA models displayed significant correlation (**Supplementary Figure 2**), with highest concordance as observed in the highest density regions (HDRs) plots (***Figure 2C***). On average, the PLS-DA models identified 1,764 features with VIPs >1 involved in subject discrimination. Notably, when considering exclusively the top 100 features obtained in the MeOH models, 86%, 85%, and 92% were also recovered and classified as significant by the EtOH models. Features of interest identified by the multivariate analysis were also investigated via univariate analysis. A high degree of correlation (R > 0.9) was observed for the feature log2 fold changes (Log2FC) between subjects obtained via the two different extraction methods (***Figure 2D***).

**Figure 2.**
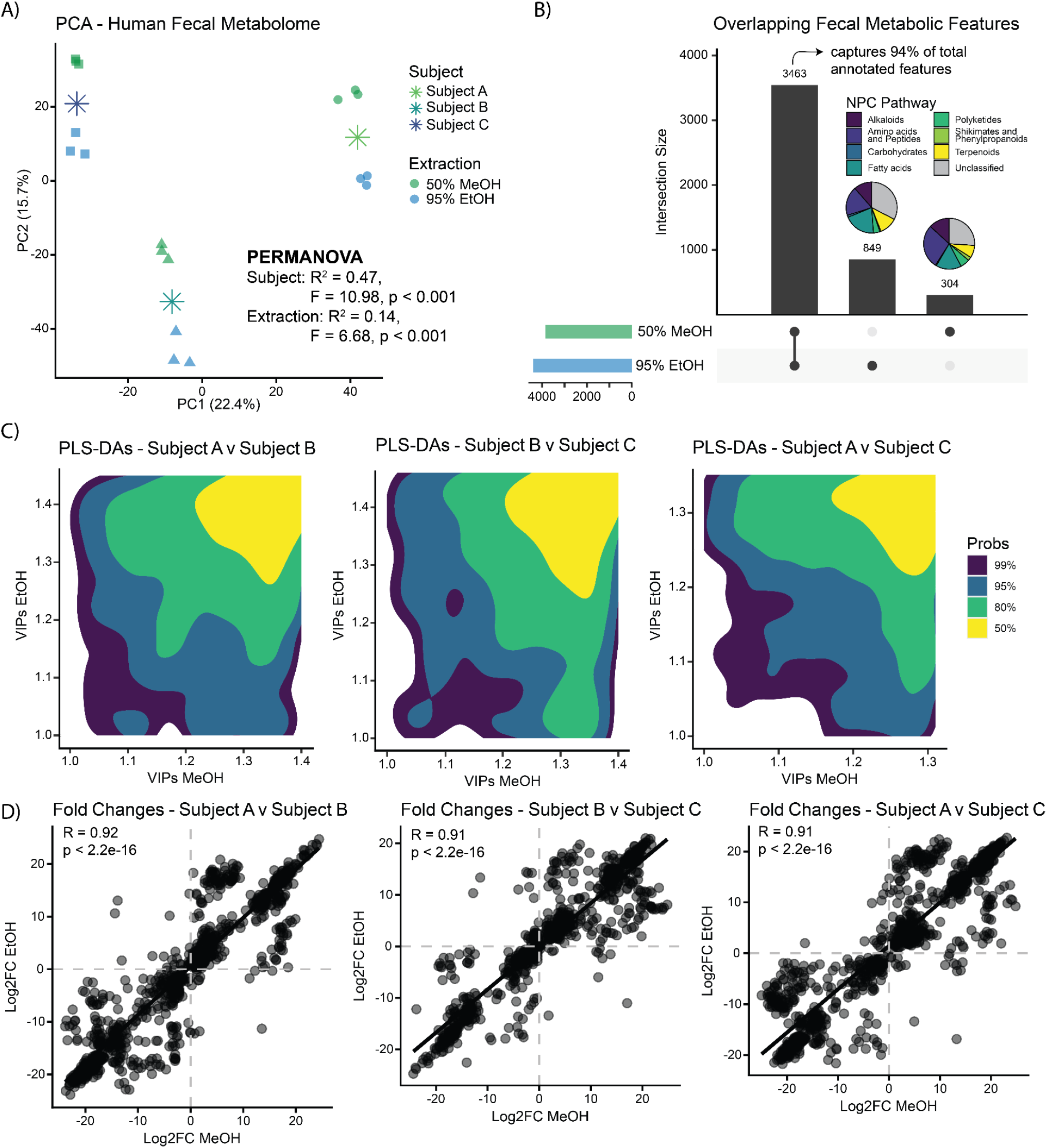
Data analysis comparison between EtOH and MeOH extractions. **A)** PCA on robust center log ratio (RCLR) transformed peak areas shows clear clustering based on subject (PERMANOVA: Subject R^2^ = 0.47, p<0.001 and Extraction R^2^ = 0.14, p<0.001). **B)** Upset plot displays that more than 75% of the obtained features are recovered by both extraction methods, encompassing 94% of the total annotated features. Additionally, EtOH appears to recover a higher number of oligopeptides, glycerolipids, and fatty amides whereas MeOH recovers more small peptides and fatty acid conjugates (***Supplementary Table 1***). **C)** HDRs plots of feature VIP scores obtained via subject pairwise PLS-DA models stratified for extraction method display a high degree of concordance. **D)** Scatter plots of the pairwise log2 fold changes (Log2FC) obtained via pairwise subject comparison in both extraction methods. Linear regression shows significant correlation (R > 0.9, p<2.2E-16) between the two methods. Asterisks indicate group centroids.

### The fecal metabolome remains stable for up to one week at RT in 95% EtOH

PCA of triplicates exclusively extracted via the 95% EtOH pipeline showed clear clustering of samples based on subject ID (PERMANOVA, R^2^ = 0.50, F = 15.24, p < 0.001) and a smaller effect of sample storage time (PERMANOVA, R^2^ = 0.08, F = 2.56, p < 0.001), which included immediate processing or storage at RT for either a day or a week in 95% EtOH (***Figure 3A***). Out of the 4,566 features obtained via EtOH extraction, 88% (3,996) were captured at all timepoints, comprising 94% (231) of all the annotated features (***Figure 3B***). The majority of the features distinctively characterizing immediate (imm), week, immediate and day, and immediate and week extractions (156 out of 570) were not classified by CANOPUS (***Supplementary Table 2***). The subsequent most affected predicted classes were small peptides (65) and oligopeptides (48). Additionally, to determine if storage time at RT could affect the metabolic features responsible for subject classification, 27 subject pairwise PLS-DA models were generated comparing samples of each subject at each different timepoint. Comparisons between immediate extractions were considered as “ground truths” and the relative recovery of those significant features was investigated for the different timepoints (***Figure 3C***). On average, 95.6%, 91.6% and 94.7% of the top 100 “ground truth” features were recovered for each pairwise subject comparison. This suggested that the fecal metabolome could remain relatively stable in 95% EtOH for up to one week at RT without losing predictive power for subject discrimination. Feature log2 fold changes across time (immediate vs 1 week) were also explored in each subject (***Supplementary Figure 3A***). Interestingly, the median fold changes of “ground truth” features obtained from the pairwise PLS-DA models were 0.705 (Subject A vs Subject B), 0.430 (Subject B vs Subject C), and 0.530 (Subject A vs Subject C) respectively, suggesting stability through time of the features of interest (***Supplementary Figure 3B***).

**Figure 3.**
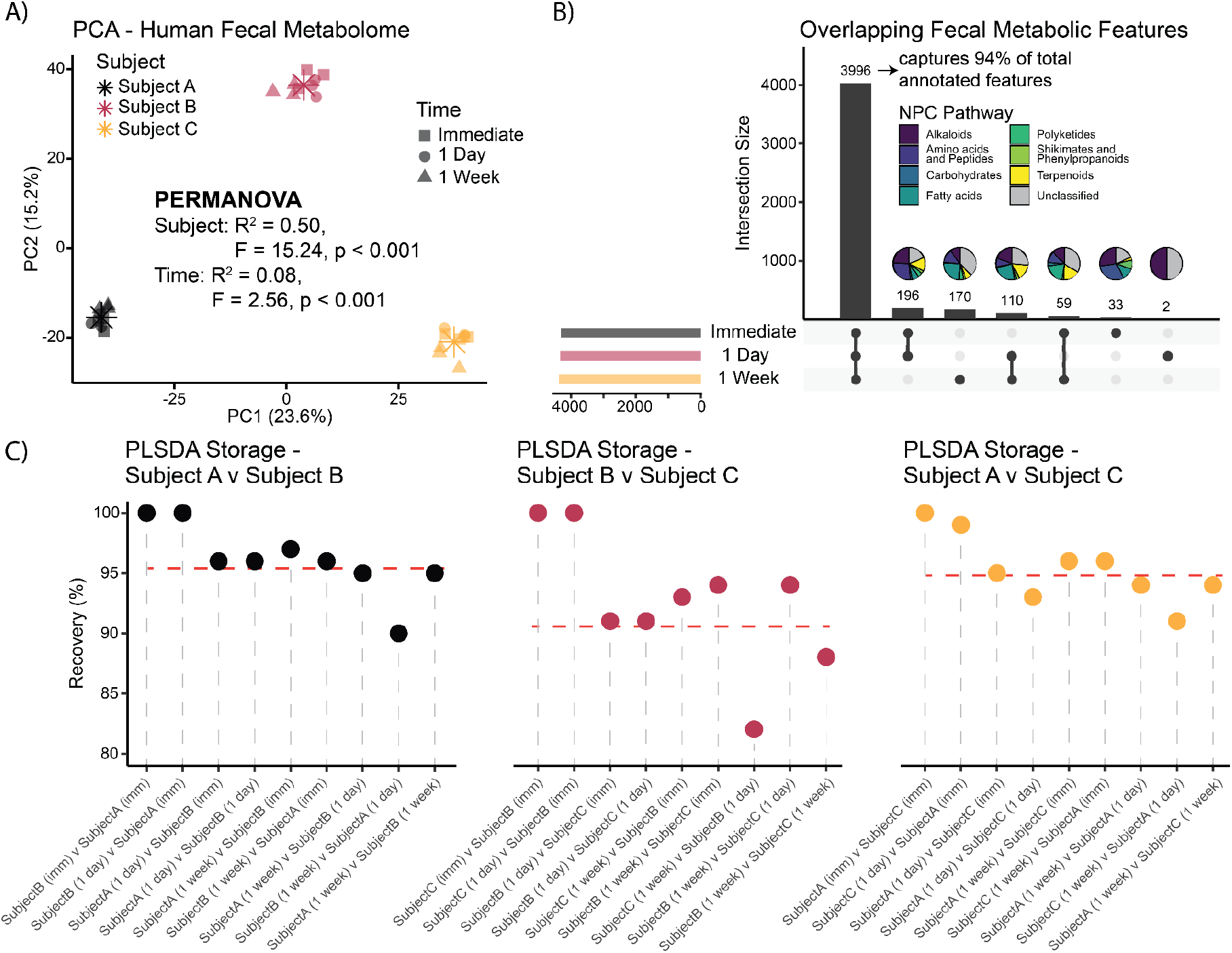
The fecal metabolome is relatively stable for up to one week at RT in EtOH. **A)** PCA on RCLR transformed peak areas shows strong clustering based on subject id and little effect of storage time (PERMANOVA: Subject R^2^ = 0.50, p<0.001 and Extraction R^2^ = 0.08, p=0.015). **B)** Upset plot displays that more than 88% of the obtained features are recovered via both extraction methods, these include 94% of the total annotated features. Most of the features exclusively characterizing the different storage timepoints are not annotated and no class prediction can be obtained via CANOPUS. **C)** Lollipop plots displaying percentage of recovery of the top 100 features discriminating subjects via pairwise PLS-DA models. Immediate extractions (imm vs imm) are considered the “ground truth” (100%). Dashed lines represent the average percentage of recovery of the “ground truth” obtained via pairwise models constructed using different storage timepoints. Asterisks indicate group centroids.

### Robust center log ratio transformation addresses sample collection discrepancies

A recurrent issue in off-site sample acquisition is the collection of fecal samples of uneven weight due to variability in water content and inconsistency in the amount collected by individuals. This can affect the metabolite recovery per sample and introduce bias in subsequent downstream analysis. Although fecal samples can be weighed before analysis, this is time consuming and technical errors can be introduced, such as repeated freeze-thaw cycles, intersample contamination, and others. To overcome these limitations, we investigated the use of RCLR transformation in untargeted metabolomics data as a method to improve the accuracy in data analysis. Originally introduced to tackle the microbiome data compositionality^29^, the RCLR transformation suits the semi-compositionality nature of the untargeted metabolomics data. Additionally, RCLR allows for easy interpretability and seamless multi-omics integration with RCLR transformed microbiome data using tools such as Joint-RPCA^31^ and DIABLO^32^. We investigated the extraction of different sample weights, 10 mg, 20 mg, and 30 mg, in both 95% EtOH and 50% MeOH extraction solvents. PCA of RCLR data shows no effect of sample weight in both extraction methods (PERMANOVAs, R^2^ < 0.05, F < 1.64, p > 0.08), with most of the variance explained by subject id and no difference in the dispersion of the samples between the different subjects (***Figure 4A***). PCA of raw or relative abundance data displayed less tightened clusters from the same subjects, significant differences between group variances (PERMDISP2, p < 0.001), and small but significant effects of sample weight (***Supplementary Figure 4***). Generated upset plot revealed that metabolite detection was not affected by sample weight, suggesting that a sample mass of 10 mg is sufficient to cover the detectable fecal metabolome in reverse phase liquid chromatography and mass spectrometry data acquired in positive ionization mode (***Figure 4B***). Finally, we explored possible correlations between weight, total obtained gDNA (genomic DNA, ng/µL), and cumulative peak areas from samples extracted with 95% EtOH and stored for one week at RT. Linear mixed effects models, accounting for repeated measures on subjects, found weight to be a significant predictor of total extracted gDNA (β = 0.25, SE = 0.08, t(23) = 2.92, p = 0.00768; ***Supplementary Figure 5A***) and the log of total extracted peak areas (β = 0.01, SE = 0.001, t(23) = 9.23, p = 3.36E-9; ***Supplementary Figure 5B***). Interestingly, gDNA appeared also to have a significant positive effect on the total extracted peaks areas (β = 0.01, SE = 0.004, t(24.98) = 2.441, p = 0.0221), but with variation in the slopes and intercepts between subjects (***Figure 4C***).

**Figure 4.**
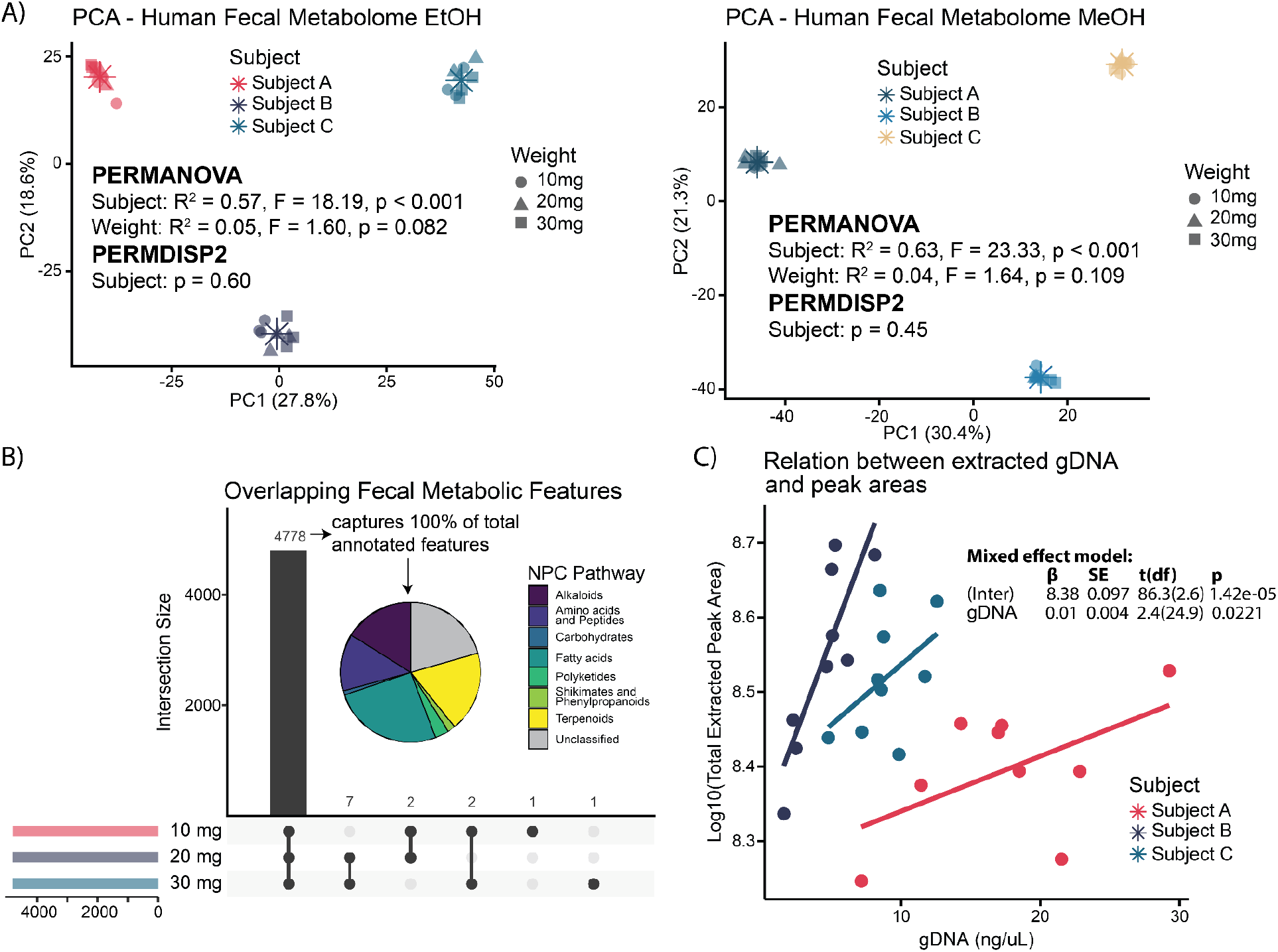
Robust center log ratio transformation in untargeted metabolomics data. **A)** PCA on RCLR transformed peak areas removes sample weight variance in both 95% EtOH and 50% MeOH extractions (PERMANOVA: Weight R^2^ < 0.05, p > 0.08) and creates homogeneous variance for samples belonging to the same subject. **B)** Upset plot displays that 99.9% of the features are captured by all the different weight aliquots, also comprising all the annotated features. Pie chart displays CANOPUS predicted NPC pathways. **C)** Scatter plot illustrates that extracted gDNA from the fecal samples is positively associated to the total extracted peak areas. Linear mixed effect model with subject id as random effect (β = 0.010, SE = 0.004, p = 0.0221). β = estimate, SE = standard error. Asterisks indicate group centroids.

## Limitations

This study focuses exclusively on human fecal samples and it has not been validated for other types of biosamples. Data was acquired via reverse phase liquid chromatography and positive ionization mode. For this reason, non detected features might display a different behavior. Provided annotations are obtained via parent ion *m/z* and MS/MS spectral matching, resulting in a level 2 annotation according to the metabolomics standards initiative^33^.

## Conclusions

The presented study highlights that the new Matrix Method, using 95% EtOH extraction, performs as well as the commonly used 50% MeOH extraction method in untargeted human fecal metabolomics studies. Moreover, EtOH is safer to use compared to MeOH and can be easily handled in off-site collections. We showcase how the fecal metabolome remains relatively stable in 95% EtOH for up to one week at RT, maintaining discriminating power between subjects. This is important as, in current approaches, samples are not always immediately stored in the most suitable condition and shipping can be costly if refrigeration is required. Finally, we highlight how data analysis via robust center log ratio transformation helps remove variance introduced by uneven sample collection and discuss how this transformation suits better integrative microbiome studies. In conclusion, 95% ethanol extraction represents a valid alternative to the more widely used 50% methanol extraction.

## Authors Contribution

S.Z. designed the study, performed MeOH extraction and data analysis, and wrote the manuscript.

V.C.L. performed MeOH extraction and ran the untargeted metabolomics experiments.

C.B. provided fecal samples, designed and performed EtOH extraction.

M.A. performed EtOH extraction.

R.K. and P.C.D. secured fundings and provided supervision. All authors reviewed and approved the manuscript.

## Supplementary Material

**Supplementary Figure 1.**
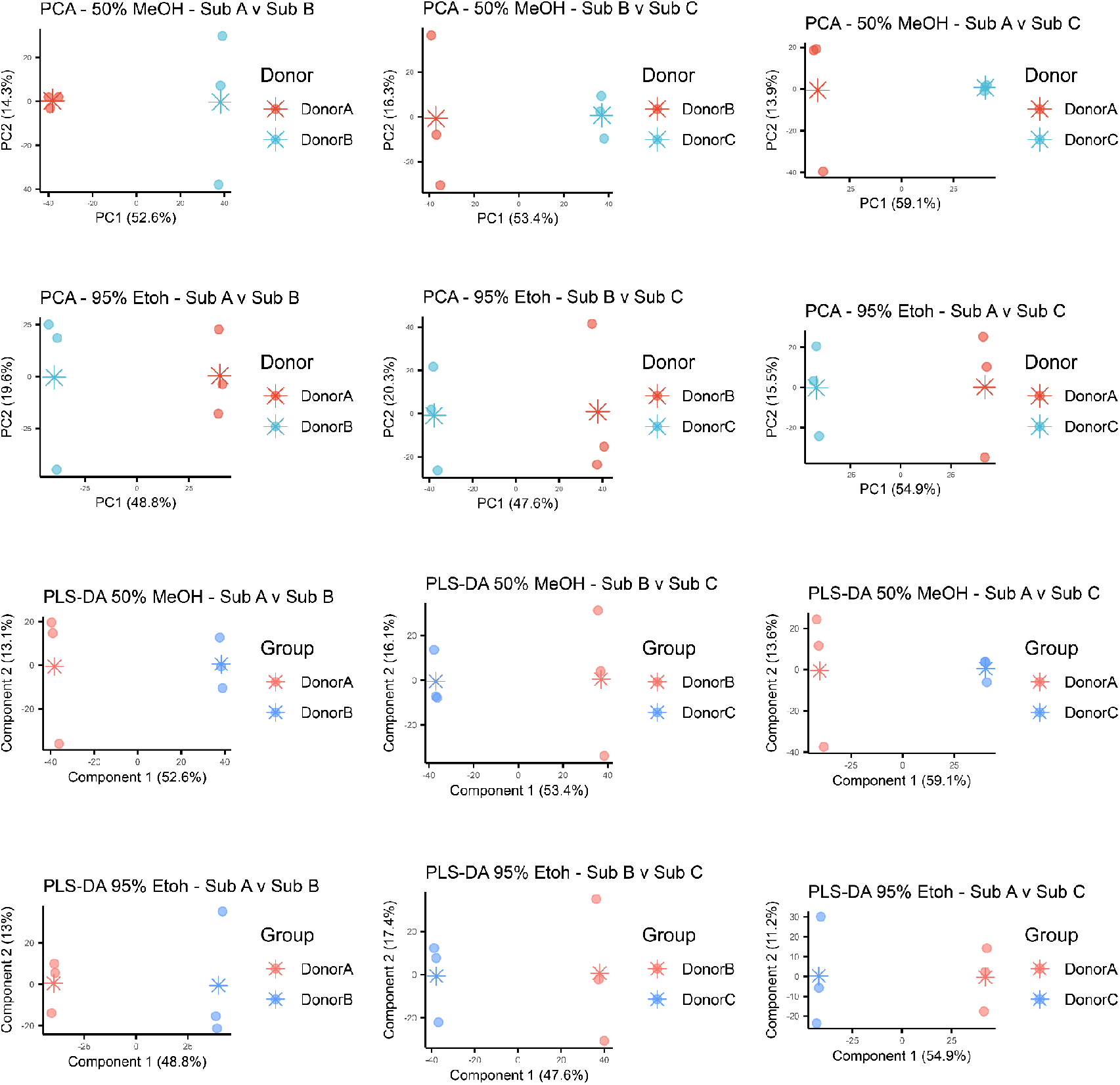
Pairwise PCA and PLS-DA models. PCA and PLS-DA score plots of the pairwise comparisons between subjects with the different extraction methods.

**Supplementary Figure 2.**
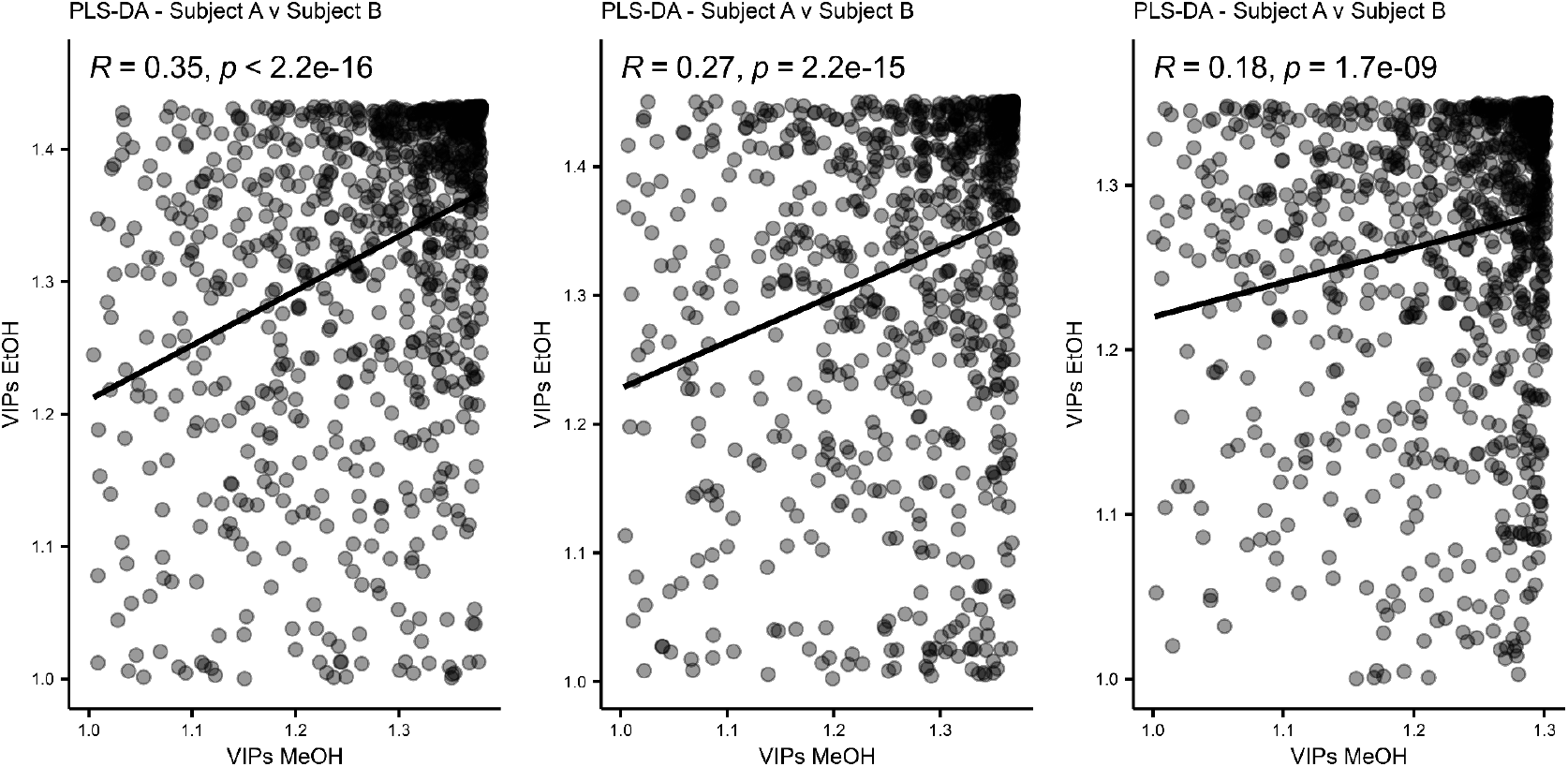
VIP score correlations. Scatter plots with linear regression analysis between VIPs of discriminating features obtained via pairwise PLS-DA models through the different extraction methods (95% EtOH and 50% MeOH).

**Supplementary Figure 3.**
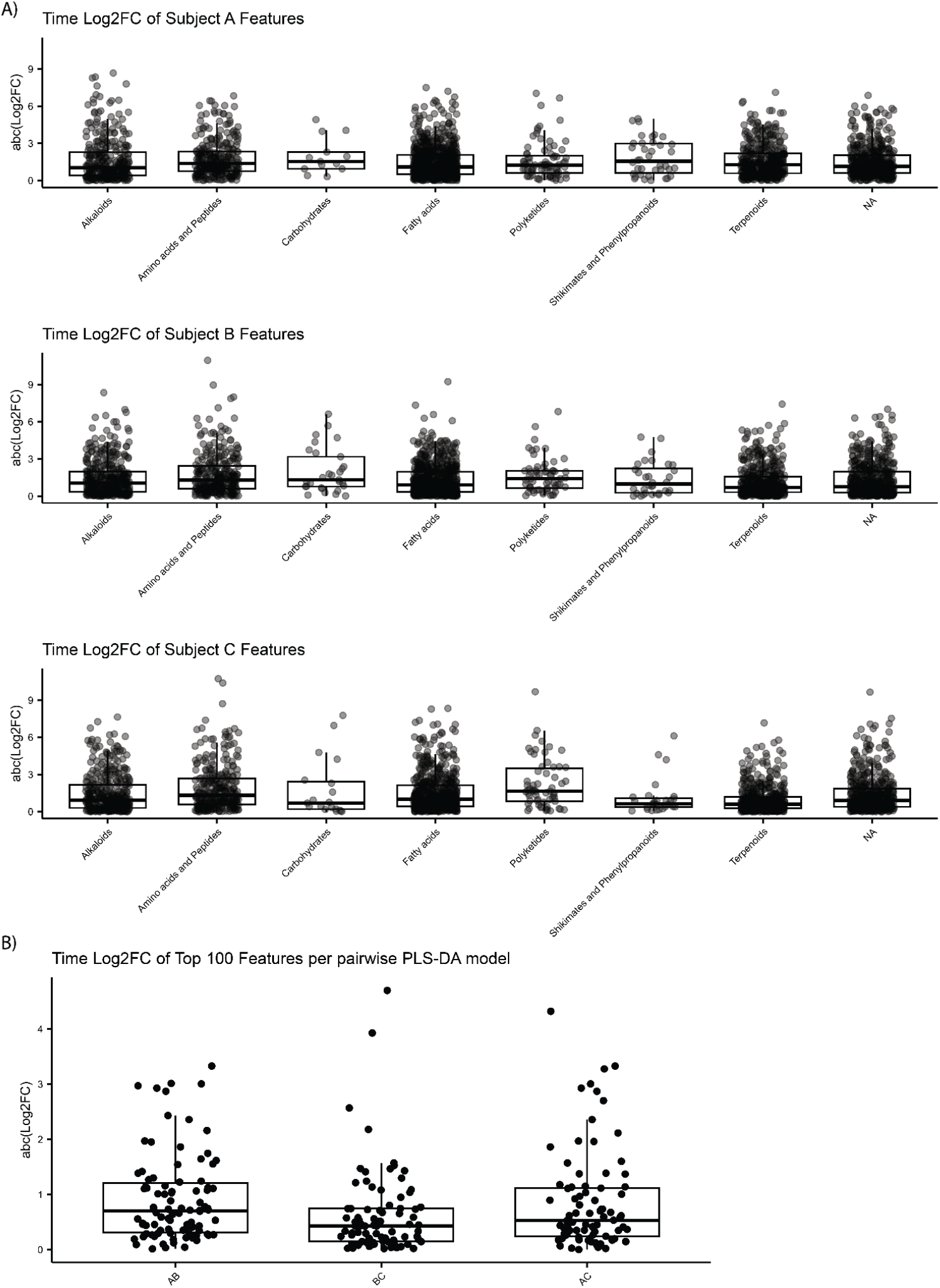
Fold changes at RT storage. **A**) Boxplots showcasing log2 fold changes (Log2FC) for each feature between immediate processing and one week storage at room temperature for the different subjects. **B**) Boxplots of the top 100 features of interest showcase a media good stability (median Log2FC < 1) of the feature of interest over a one week storage at room temperature.

**Supplementary Figure 4.**
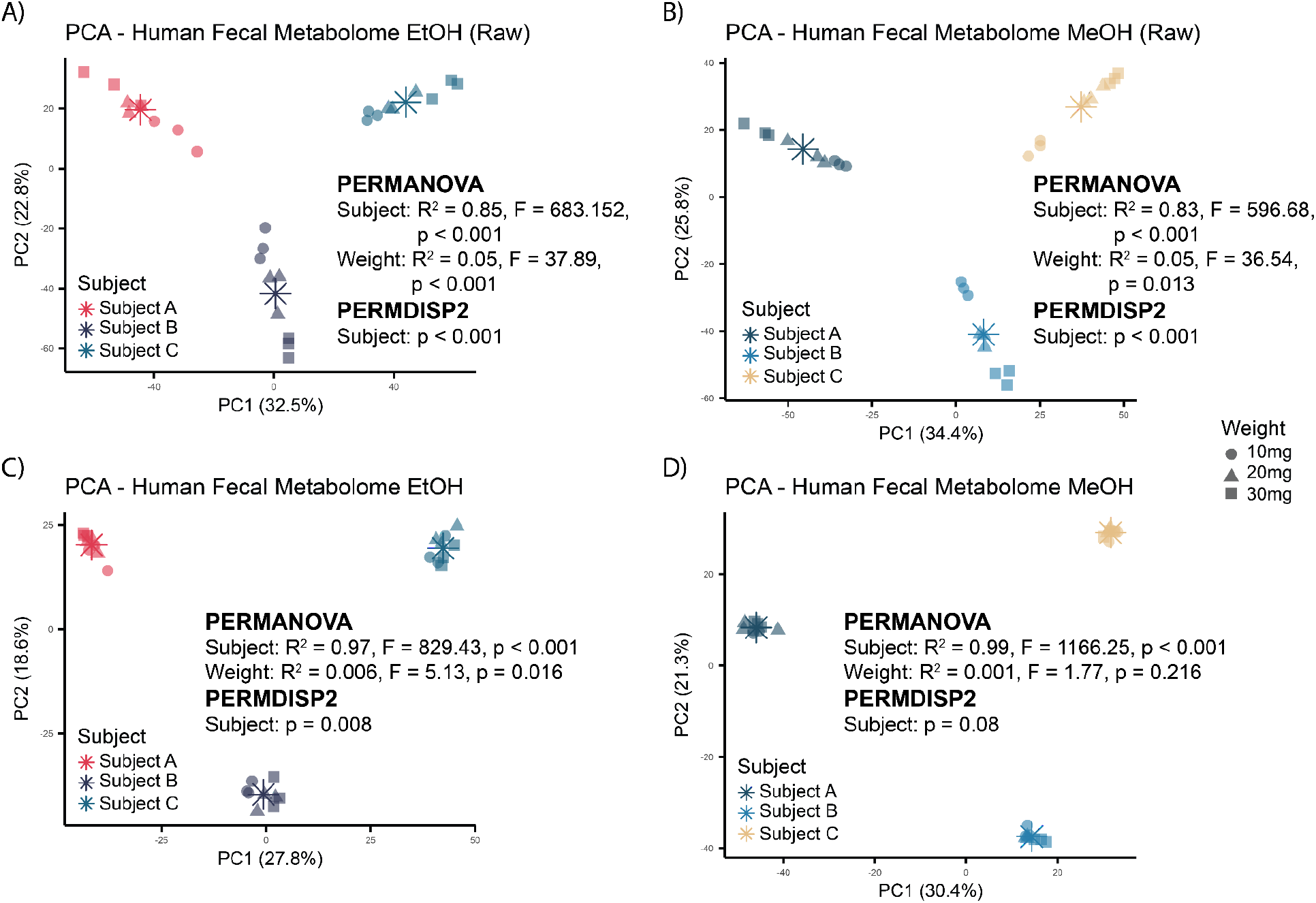
Data transformation effects. PCA score plots of aliquots of different weights (10, 20, and 30 mg). PCA on raw data for both EtOH and MeOH extractions (**A and B**) displayed variance based on sample weight (PERMANOVA, R^2^ = 0.05, p < 0.05) and differences in variance homogeneity between subjects (PERMDISP2, p < 0.001). Relative abundance normalization (**C and D**), also known as TIC normalization, partially reduces weight effect (PERMANOVA, R^2^ < 0.008, p = 0.016 and 0.216) but does not completely uniform variance between subjects (PERMDISP2, p = 0.008 and p = 0.08).

**Supplementary Figure 5.**
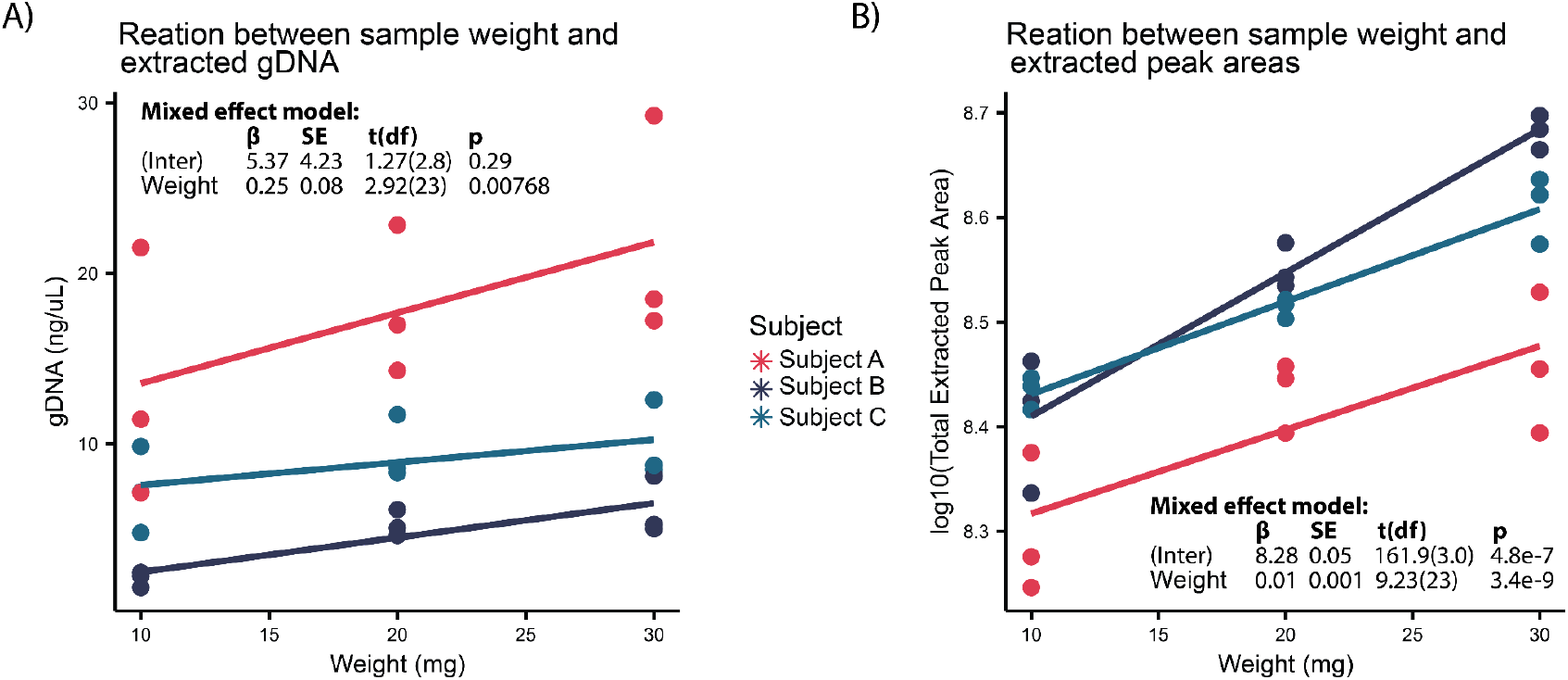
Correlation between sample weight and extracted gDNA or TIC. Scatter plots showcase positive correlation between weight of the samples and total extracted gDNA (**A**) and peak areas (**B**). Repeated measures of the same subject were taken into account via linear mixed effect models.

**Supplementary Table 1.**
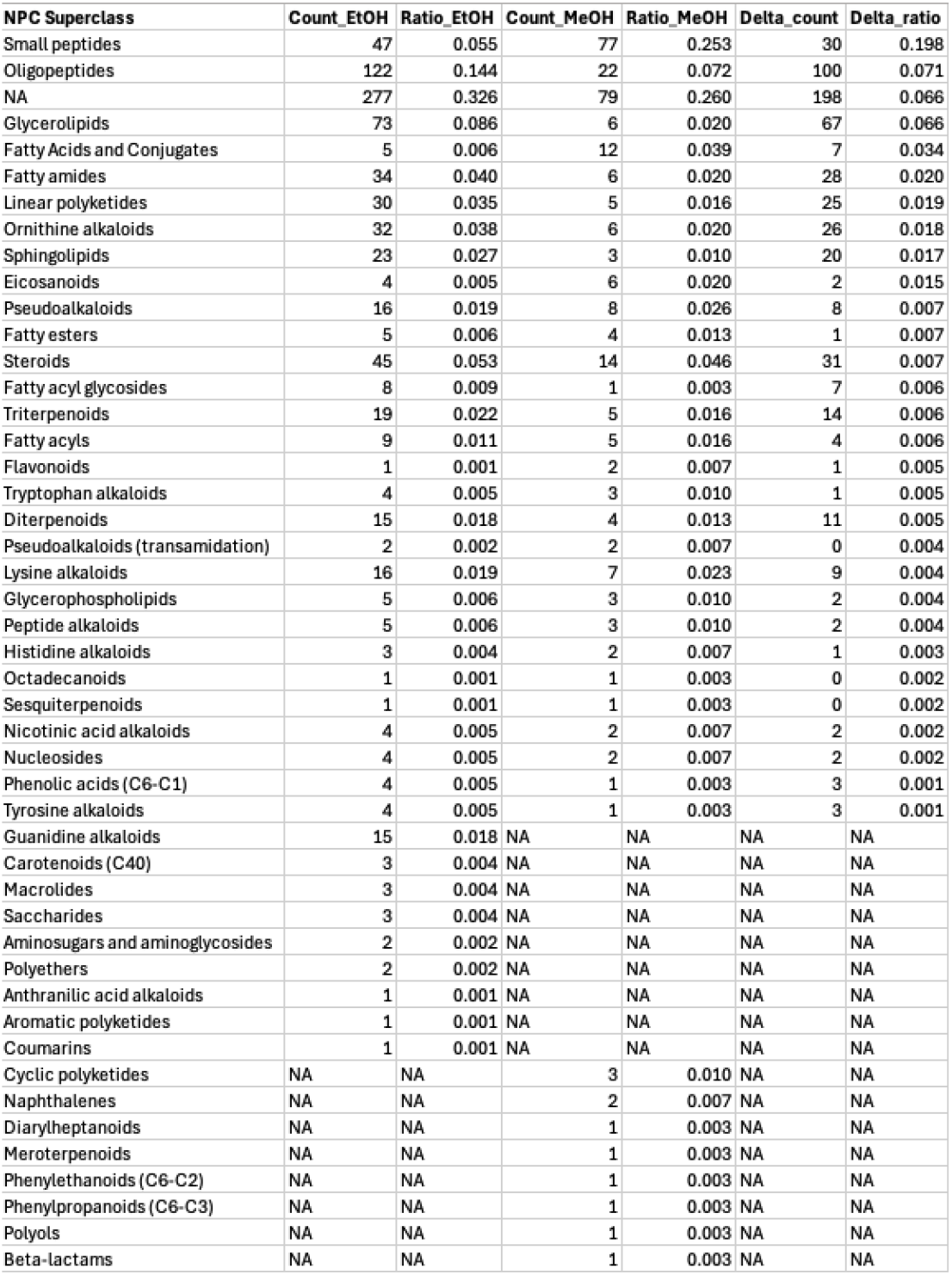
Extraction specific (MeOH v EtOH) feature class prediction.

| NPC Superclass | Count_EtOH | Ratio_EtOH | Count_MeOH | Ratio_MeOH | Delta_count | Delta_ratio |
| --- | --- | --- | --- | --- | --- | --- |
| Small peptides | 47 | 0.055 | 77 | 0.253 | 30 | 0.198 |
| Oligopeptides | 122 | 0.144 | 22 | 0.072 | 100 | 0.071 |
| NA | 277 | 0.326 | 79 | 0.260 | 198 | 0.066 |
| Glycerolipids | 73 | 0.086 | 6 | 0.020 | 67 | 0.066 |
| Fatty Acids and Conjugates | 5 | 0.006 | 12 | 0.039 | 7 | 0.034 |
| Fatty amides | 34 | 0.040 | 6 | 0.020 | 28 | 0.020 |
| Linear polyketides | 30 | 0.035 | 5 | 0.016 | 25 | 0.019 |
| Ornithine alkaloids | 32 | 0.038 | 6 | 0.020 | 26 | 0.018 |
| Sphingolipids | 23 | 0.027 | 3 | 0.010 | 20 | 0.017 |
| Eicosanoids | 4 | 0.005 | 6 | 0.020 | 2 | 0.015 |
| Pseudoalkaloids | 16 | 0.019 | 8 | 0.026 | 8 | 0.007 |
| Fatty esters | 5 | 0.006 | 4 | 0.013 | 1 | 0.007 |
| Steroids | 45 | 0.053 | 14 | 0.046 | 31 | 0.007 |
| Fatty acyl glycosides | 8 | 0.009 | 1 | 0.003 | 7 | 0.006 |
| Triterpenoids | 19 | 0.022 | 5 | 0.016 | 14 | 0.006 |
| Fatty acyls | 9 | 0.011 | 5 | 0.016 | 4 | 0.006 |
| Flavonoids | 1 | 0.001 | 2 | 0.007 | 1 | 0.005 |
| Tryptophan alkaloids | 4 | 0.005 | 3 | 0.010 | 1 | 0.005 |
| Diterpenoids | 15 | 0.018 | 4 | 0.013 | 11 | 0.005 |
| Pseudoalkaloids (transamidation) | 2 | 0.002 | 2 | 0.007 | 0 | 0.004 |
| Lysine alkaloids | 16 | 0.019 | 7 | 0.023 | 9 | 0.004 |
| Glycerophospholipids | 5 | 0.006 | 3 | 0.010 | 2 | 0.004 |
| Peptide alkaloids | 5 | 0.006 | 3 | 0.010 | 2 | 0.004 |
| Histidine alkaloids | 3 | 0.004 | 2 | 0.007 | 1 | 0.003 |
| Octadecanoids | 1 | 0.001 | 1 | 0.003 | 0 | 0.002 |
| Sesquiterpenoids | 1 | 0.001 | 1 | 0.003 | 0 | 0.002 |
| Nicotinic acid alkaloids | 4 | 0.005 | 2 | 0.007 | 2 | 0.002 |
| Nucleosides | 4 | 0.005 | 2 | 0.007 | 2 | 0.002 |
| Phenolic acids (C6-C1) | 4 | 0.005 | 1 | 0.003 | 3 | 0.001 |
| Tyrosine alkaloids | 4 | 0.005 | 1 | 0.003 | 3 | 0.001 |
| Guanidine alkaloids | 15 | 0.018 | NA | NA | NA | NA |
| Carotenoids (C40) | 3 | 0.004 | NA | NA | NA | NA |
| Macrolides | 3 | 0.004 | NA | NA | NA | NA |
| Saccharides | 3 | 0.004 | NA | NA | NA | NA |
| Aminosugars and aminoglycosides | 2 | 0.002 | NA | NA | NA | NA |
| Polyethers | 2 | 0.002 | NA | NA | NA | NA |
| Anthranilic acid alkaloids | 1 | 0.001 | NA | NA | NA | NA |
| Aromatic polyketides | 1 | 0.001 | NA | NA | NA | NA |
| Coumarins | 1 | 0.001 | NA | NA | NA | NA |
| Cyclic polyketides | NA | NA | 3 | 0.010 | NA | NA |
| Naphthalenes | NA | NA | 2 | 0.007 | NA | NA |
| Diarylheptanoids | NA | NA | 1 | 0.003 | NA | NA |
| Meroterpenoids | NA | NA | 1 | 0.003 | NA | NA |
| Phenylethanoids (C6-C2) | NA | NA | 1 | 0.003 | NA | NA |
| Phenylpropanoids (C6-C3) | NA | NA | 1 | 0.003 | NA | NA |
| Polyols | NA | NA | 1 | 0.003 | NA | NA |
| Beta-lactams | NA | NA | 1 | 0.003 | NA | NA |

**Supplementary Table 2.** Extraction specific (storage) feature class prediction.

| NPC Superclass | Count | Ratio | Time | NPC Superclass | Count | Ratio | Time | NPC Superclass | Count | Ratio | Time |
| --- | --- | --- | --- | --- | --- | --- | --- | --- | --- | --- | --- |
| Aminosugars and aminoglycosides | 1 | 0.005 | Immediate_day | Linear polyketides | 1 | 0.005 | Immediate_day | Phenolic acids (C6-C1) | 2 | 0.012 | Week_exclusive |
| Aminosugars and aminoglycosides | 1 | 0.006 | Week_exclusive | Linear polyketides | 2 | 0.018 | Day_week | Phenolic acids (C6-C1) | 1 | 0.030 | Immediate_exclusive |
| Aminosugars and aminoglycosides | 1 | 0.017 | Immediate_week | Linear polyketides | 1 | 0.030 | Immediate_exclusive | Phenylpropanoids (C6-C3) | 1 | 0.006 | Week_exclusive |
| Anthranilic acid alkaloids | 1 | 0.005 | Immediate_day | Linear polyketides | 6 | 0.035 | Week_exclusive | Phenylpropanoids (C6-C3) | 3 | 0.015 | Immediate_day |
| Anthranilic acid alkaloids | 1 | 0.017 | Immediate_week | Linear polyketides | 3 | 0.051 | Immediate_week | Pseudoalkaloids | 1 | 0.006 | Week_exclusive |
| Anthranilic acid alkaloids | 1 | 0.030 | Immediate_exclusive | Lysine alkaloids | 3 | 0.018 | Week_exclusive | Pseudoalkaloids | 3 | 0.015 | Immediate_day |
| Coumarins | 1 | 0.005 | Immediate_day | Lysine alkaloids | 3 | 0.027 | Day_week | Pseudoalkaloids | 1 | 0.017 | Immediate_week |
| Cyclic polyketides | 1 | 0.005 | Immediate_day | Lysine alkaloids | 2 | 0.034 | Immediate_week | Pseudoalkaloids | 1 | 0.030 | Immediate_exclusive |
| Diarylheptanoids | 1 | 0.006 | Week_exclusive | Lysine alkaloids | 2 | 0.061 | Immediate_exclusive | Pseudoalkaloids | 6 | 0.055 | Day_week |
| Diterpenoids | 2 | 0.018 | Day_week | Lysine alkaloids | 13 | 0.066 | Immediate_day | Pseudoalkaloids (transamidation) | 1 | 0.006 | Week_exclusive |
| Diterpenoids | 1 | 0.030 | Immediate_exclusive | Macrolides | 2 | 0.010 | Immediate_day | Pseudoalkaloids (transamidation) | 2 | 0.018 | Day_week |
| Diterpenoids | 6 | 0.031 | Immediate_day | Macrolides | 1 | 0.017 | Immediate_week | Saccharides | 1 | 0.005 | Immediate_day |
| Diterpenoids | 6 | 0.035 | Week_exclusive | Macrolides | 3 | 0.018 | Week_exclusive | Sesquiterpenoids | 1 | 0.005 | Immediate_day |
| Diterpenoids | 4 | 0.068 | Immediate_week | Meroterpenoids | 1 | 0.006 | Week_exclusive | Sesquiterpenoids | 1 | 0.006 | Week_exclusive |
| Eicosanoids | 2 | 0.012 | Week_exclusive | Meroterpenoids | 2 | 0.010 | Immediate_day | Small peptides | 6 | 0.035 | Week_exclusive |
| Eicosanoids | 3 | 0.027 | Day_week | Meroterpenoids | 1 | 0.030 | Immediate_exclusive | Small peptides | 8 | 0.073 | Day_week |
| Fatty Acids and Conjugates | 3 | 0.015 | Immediate_day | NA | 6 | 0.182 | Immediate_exclusive | Small peptides | 6 | 0.102 | Immediate_week |
| Fatty Acids and Conjugates | 5 | 0.029 | Week_exclusive | NA | 36 | 0.184 | Immediate_day | Small peptides | 37 | 0.189 | Immediate_day |
| Fatty Acids and Conjugates | 1 | 0.030 | Immediate_exclusive | NA | 29 | 0.264 | Day_week | Small peptides | 7 | 0.212 | Immediate_exclusive |
| Fatty Acids and Conjugates | 2 | 0.034 | Immediate_week | NA | 20 | 0.339 | Immediate_week | Small peptides | 1 | 0.500 | Day_exclusive |
| Fatty Acids and Conjugates | 4 | 0.036 | Day_week | NA | 64 | 0.376 | Week_exclusive | Sphingolipids | 1 | 0.005 | Immediate_day |
| Fatty acyl glycosides | 1 | 0.017 | Immediate_week | NA | 1 | 0.500 | Day_exclusive | Sphingolipids | 7 | 0.041 | Week_exclusive |
| Fatty acyls | 1 | 0.006 | Week_exclusive | Nicotinic acid alkaloids | 3 | 0.015 | Immediate_day | Steroids | 5 | 0.029 | Week_exclusive |
| Fatty acyls | 3 | 0.027 | Day_week | Nicotinic acid alkaloids | 1 | 0.017 | Immediate_week | Steroids | 12 | 0.061 | Immediate_day |
| Fatty acyls | 1 | 0.030 | Immediate_exclusive | Nicotinic acid alkaloids | 2 | 0.018 | Day_week | Steroids | 8 | 0.073 | Day_week |
| Fatty amides | 3 | 0.015 | Immediate_day | Nicotinic acid alkaloids | 1 | 0.030 | Immediate_exclusive | Steroids | 5 | 0.085 | Immediate_week |
| Fatty amides | 1 | 0.030 | Immediate_exclusive | Nucleosides | 1 | 0.006 | Week_exclusive | Triterpenoids | 1 | 0.006 | Week_exclusive |
| Fatty amides | 4 | 0.036 | Day_week | Nucleosides | 1 | 0.017 | Immediate_week | Triterpenoids | 4 | 0.020 | Immediate_day |
| Fatty amides | 12 | 0.071 | Week_exclusive | Nucleosides | 2 | 0.018 | Day_week | Triterpenoids | 4 | 0.036 | Day_week |
| Fatty amides | 5 | 0.085 | Immediate_week | Octadecanoids | 1 | 0.009 | Day_week | Tryptophan alkaloids | 1 | 0.006 | Week_exclusive |
| Fatty esters | 1 | 0.005 | Immediate_day | Octadecanoids | 1 | 0.030 | Immediate_exclusive | Tryptophan alkaloids | 1 | 0.009 | Day_week |
| Fatty esters | 2 | 0.012 | Week_exclusive | Oligopeptides | 3 | 0.027 | Day_week | Tryptophan alkaloids | 10 | 0.051 | Immediate_day |
| Fatty esters | 8 | 0.073 | Day_week | Oligopeptides | 2 | 0.034 | Immediate_week | Tyrosine alkaloids | 1 | 0.006 | Week_exclusive |
| Flavonoids | 1 | 0.005 | Immediate_day | Oligopeptides | 15 | 0.088 | Week_exclusive | Tyrosine alkaloids | 2 | 0.018 | Day_week |
| Flavonoids | 3 | 0.027 | Day_week | Oligopeptides | 4 | 0.121 | Immediate_exclusive | Xanthones | 1 | 0.005 | Immediate_day |
| Glycerolipids | 1 | 0.005 | Immediate_day | Oligopeptides | 24 | 0.122 | Immediate_day |  |  |  |  |
| Glycerolipids | 2 | 0.034 | Immediate_week | Ornithine alkaloids | 1 | 0.017 | Immediate_week |  |  |  |  |
| Glycerolipids | 4 | 0.036 | Day_week | Ornithine alkaloids | 3 | 0.027 | Day_week |  |  |  |  |
| Glycerolipids | 12 | 0.071 | Week_exclusive | Ornithine alkaloids | 5 | 0.029 | Week_exclusive |  |  |  |  |
| Glycerophospholipids | 1 | 0.005 | Immediate_day | Ornithine alkaloids | 1 | 0.030 | Immediate_exclusive |  |  |  |  |
| Glycerophospholipids | 2 | 0.018 | Day_week | Ornithine alkaloids | 12 | 0.061 | Immediate_day |  |  |  |  |
| Histidine alkaloids | 1 | 0.006 | Week_exclusive | Peptide alkaloids | 1 | 0.009 | Day_week |  |  |  |  |
| Histidine alkaloids | 6 | 0.031 | Immediate_day | Peptide alkaloids | 2 | 0.012 | Week_exclusive |  |  |  |  |
| Lignans | 1 | 0.005 | Immediate_day | Peptide alkaloids | 3 | 0.015 | Immediate_day |  |  |  |  |
|  |  |  |  | Peptide alkaloids | 2 | 0.061 | Immediate_exclusive |  |  |  |  |

